# Simultaneous decoding of cardiovascular and respiratory functional changes from pig intraneural vagus nerve signals

**DOI:** 10.1101/2020.06.01.127050

**Authors:** Fabio Vallone, Matteo Maria Ottaviani, Francesca Dedola, Annarita Cutrone, Simone Romeni, Adele Macrí Panarese, Fabio Bernini, Marina Cracchiolo, Khatia Gabisonia, Nikoloz Gorgodze, Alberto Mazzoni, Fabio A. Recchia, Silvestro Micera

## Abstract

Bioelectronic medicine is opening new perspectives for the treatment of some major chronic diseases through the physical modulation of autonomic nervous system activity. Being the main peripheral route for electrical signals between central nervous system and visceral organs, the vagus nerve (VN) is one of the most promising targets. Closed-loop neuromodulation would be crucial to increase effectiveness and reduce side effects, but it depends on the possibility of extracting useful physiological information from VN electrical activity, which is currently very limited.

Here, we present a new decoding algorithm properly detecting different functional changes from VN signals. They were recorded using intraneural electrodes in anaesthetized pigs during cardiovascular and respiratory challenges mimicking increases in arterial blood pressure, tidal volume and respiratory rate. A novel decoding algorithm was developed combining discrete wavelet transformation, principal component analysis, and ensemble learning made of classification trees. It robustly achieved high accuracy levels in identifying different functional changes and discriminating among them. We also introduced a new index for the characterization of recording and decoding performance of neural interfaces. Finally, by combining an anatomically validated hybrid neural model and discrimination analysis, we provided new evidence suggesting a functional topographical organization of VN fascicles. This study represents an important step towards the comprehension of VN signaling, paving the way to the development of effective closed-loop bioelectronic systems.

## Introduction

The autonomic nervous system (ANS) plays a crucial role in the self-governed maintenance of body homeostasis. In ANS peripheral nerves, afferent and efferent fibres run together, providing bidirectional communication between specific circuits of the central nervous system and visceral organs. The artificial modulation of this complex circuitry is the challenging goal of bioelectronic medicine (BM), a highly promising alternative to some limited pharmacological tretments^1–3^. Among the main ANS nerves, the vagus nerve (VN) represents a privileged target as it modulates vital functions like respiration, circulation and the digestion^4^. VN stimulation (VNS) of cervical segments has shown a great potential for the treatment of a wide range of pathological conditions such as epilepsy^5^, chronic heart failure^6^, and inflammatory diseases^7,8^. However, the formidable amount of afferent and efferent signals that simultaneously cross this VN segment, the numerous VNS side-effects^9^ and the discovery of VN involvement in the regulation of complex functions like immunity^10^ or central neuroplasticity^11,12^ highlight the need for high precision and selectivity. In an ideal scenario, the therapeutic stimulation or inhibition of VN or any other ANS nerve should be: a) selectively directed to specific efferent or afferent fibres and b) regulated by a closed-loop feedback, thus adapting the stimulation to patient-specific conditions^13–16^. Importantly, the co-existence of afferent and efferent signals in the VN points to the possibility that the feedback loop is originated and closed at the same anatomical site. Nonetheless, VNS is currently applied in an open-loop fashion^17,18^ and mainly delivered using epineural cuff-like electrodes, because of their relatively low invasiveness and versatility for chronic applications^19,20^.

The first step for the development of a closed loop modality would be the precise identification of physiological “states” by processing the autonomic neural signals. Different strategies have been recently adopted to extract function-specific markers from neural activity using epineural electrodes^3,21–25^. Specific neurograms related to the respiratory cycle^21,22^ and blood pressure fluctuations^3^ were recorded from pig VN using bipolar/tripolar ring cuff electrodes, while decoding strategies and methodological frameworks led to the identification of cytokine- and hypoglycemia/hyperglycemia-specific neural activity markers in murine VN and carotid sinus nerve^23–25^. However, epineural electrodes display a limited selectivity and can only record the compound activity that provides a global picture of neural signal trafficking^26–28^. Therefore, intraneural electrodes have been developed and successfully employed by us and others to enhance selectivity and increase the signal-to-noise ratio of recordings^26,29^ in somatic nerves^13,16,30,31^. Nevertheless, this technology has received very limited attention for VN applications and only one preliminary study has been published to date^32^, which was based on a simple experimental protocol (steady neural activity) and a limited signal recording capacity (i.e. 4-channel electrode longitudinally implanted).

Here, we performed the first comprehensive study testing intraneural VN recording to identify physiological (arterial and respiratory) changes. To this aim, we recorded VN activity through intraneural multi-channel electrodes in anaesthetized pigs at baseline and during alterations in blood pressure and respiratory parameters, collectively named “functional challenges”. To discriminate between baseline condition and functional challenges and to investigate a possible functional organization of VN fascicles, we developed novel decoding algorithms and hybrid modeling framework (see next section and Methods).

## Results

### Experimental setup and decoding method to classify functional challenges against baseline

To study the feasibility of decoding relevant physiological states based on VN neural activity, intrafascicular multi-channel electrodes^40^ were implanted into the cervical VN of 4 anaesthetized and artificially ventilated pigs (p1,…, p4) (Fig.1 a,b). Neural activity was recorded at baseline and during functional challenges obtained by infusing the vasopressor angiotensin II (AngII) to increase baseline blood pressure (BP) by 150% and/or by varying the ventilator parameters to increase respiratory rate (RR) and/or tidal volume (TV). These 3 functional challenges, named BPC, RRC and TVC, were compared to baseline condition and among them (Fig. 1c and Methods for details). To obtain the desired vasopressor effect, AngII infusion was maintained for 8.8 ± 0.8 min (n=5) and mean BP increased from 76 ± 2.2 mmHg up to 110 ± 2.6 mmHg (n=5).

**Fig. 1.**
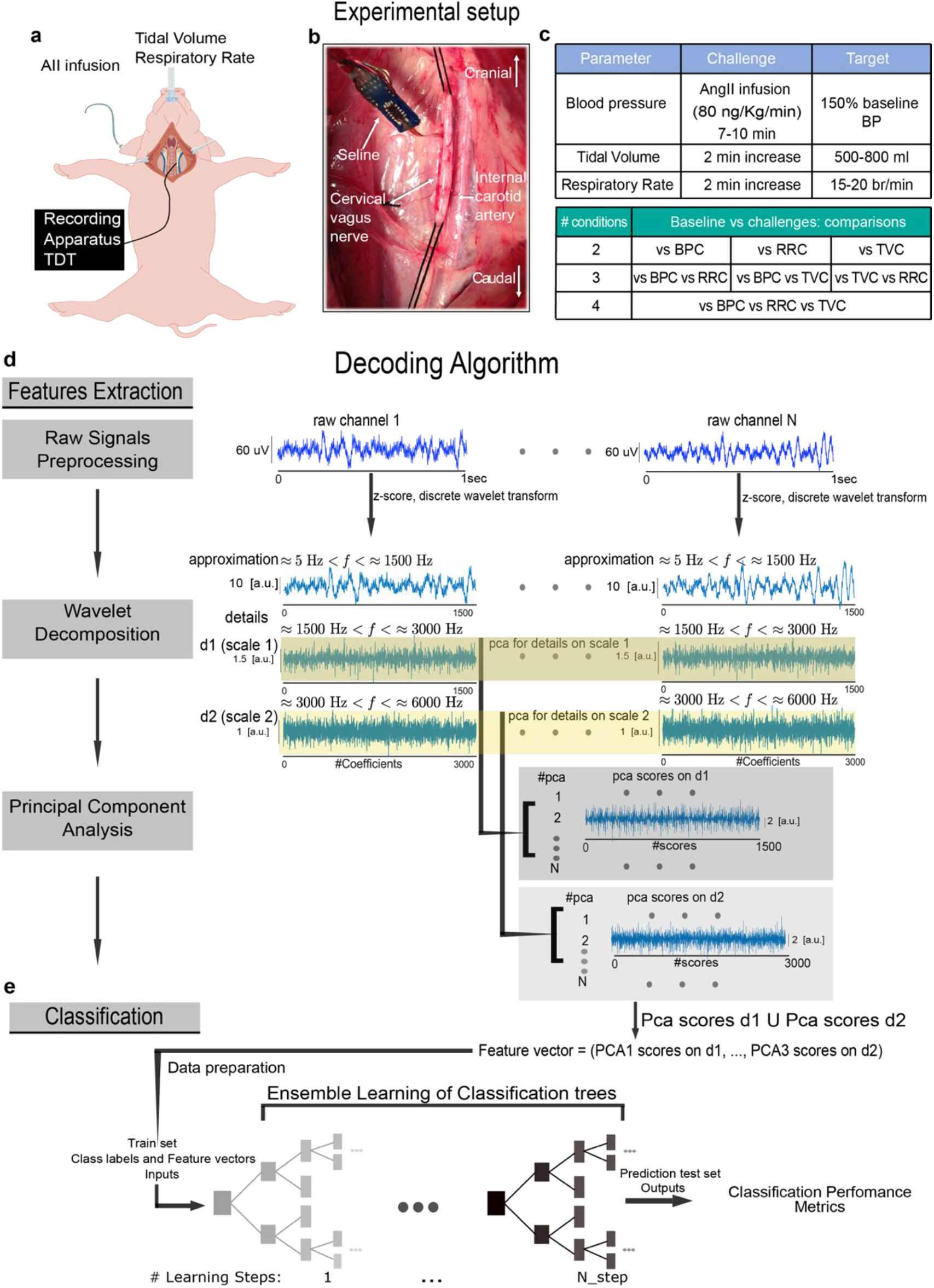
Schematic representation of the experimental setup and decoding algorithm **a,b** Recording apparatus and electrode implantation **c** Summary of the in vivo protocol and comparisons. **d** Decoding algorithm. Feature extraction was performed on raw signals by applying principal component analysis on wavelet details relative to two different scales. **e** Decoding performed with ensemble learning based on classification tree combined with random undersampling and boosting procedure (see Methods) was applied on feature vectors. Decoding performance was assessed by means of confusion matrices and accuracy level.

A decoding algorithm was developed to process multivariate signals acquired from the multichannel intraneural electrodes. To extract from the neural activity relevant features encoding for the functional challenges, we first employed a multi-scale decomposition analysis based on the discrete wavelet transform^33,34^ to extract the high frequency component (>1500 Hz) of our recordings. Subsequently, we focused on the direction of greatest variance by using principal components analysis^35^ (Fig. 1d and Methods for details). Then, an ensemble of classifiers made on classification trees^36,37^ was trained and we assessed the classification performance on the test set using accuracy values (Fig. 1e and Methods for details).

We tested our decoding algorithm in different scenarios, i.e., considering a progressively larger number and type of functional challenges. In two pigs (p3 and p4) we also studied the variation of the decoding performances relative to the positioning of two SELINEs (s1, s2) and (s3, s4) within the same nerves, thus testing the following combinations: p3-s1, p3-s2 and p4-s3, p4-s4.

### Decoding the response to a single functional challenge

We started with the simplest case in which we sought to discriminate only one of the three functional challenges against baseline. For baseline vs BPC, we achieved a mean accuracy level over the different recordings equal to 90.7±5.7% as shown in Fig. 2a (n=5, see Supplementary Fig. 1a for confusion matrices).

**Fig. 2.**
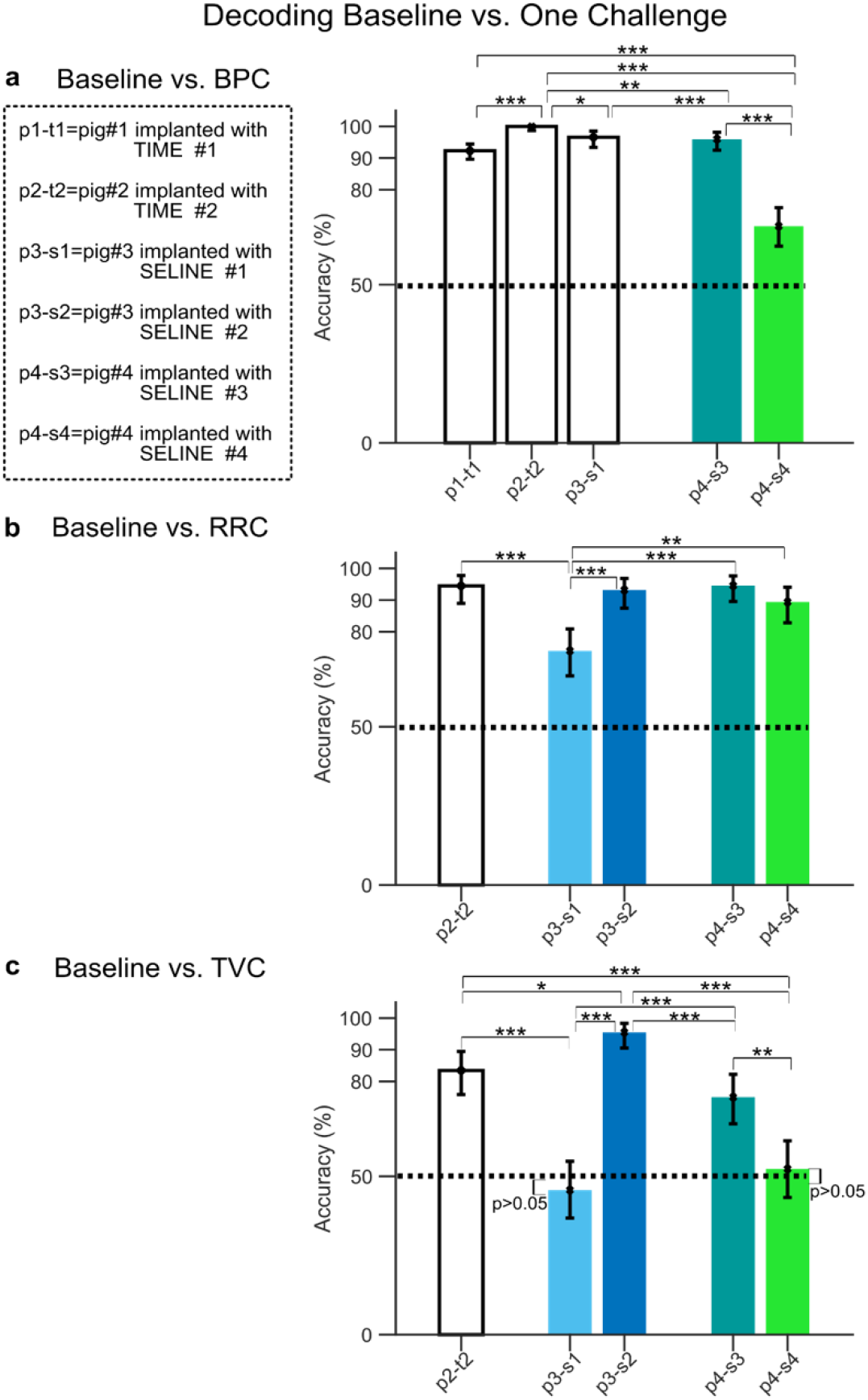
Decoding accuracy for baseline condition vs. a single functional challenge **a,b,c** Accuracy level together with p=0.05 confidence interval (error bars, Clopper-Pearson method) for the comparison between baseline and BPC (panel a), RRC (panel b) and TVC (panel c). For all panels, colored bars indicate the same animal implanted with two different electrodes. Dashed lines represent chance level. To test statistically significant differences with respect to chance level in the cases where confidence intervals overlapped with chance level, a binomial test was used (panel c, animals p3-s1 and p4-s3, p>0.05 binomial test). Statistical comparisons between animals were assessed by using a Chi square test Bonferroni corrected for multiple comparisons (*p<0.05, **p<0.01, ***p<0.001).

High-level accuracy during increased RRC was also obtained, equal to 89.1 ±3.9%, as shown in Fig. 2b (n=4, see also Supplementary Fig. 1b for confusion matrices). Measurements during TVC yielded more heterogeneous results compared to the other challenges. In pig p2 that was implanted with TIME t2 (p2-t2), we achieved high accuracy level equal to 83.5% ([75.8%, 89.5%], Confidence Interval p=0.05, see Fig. 2c and Supplementary Fig. 1c for confusion matrices).

Moreover, the decoding accuracy during TVC was strongly dependent on the electrode positioning. In fact, in animals implanted with s1 and s4, i.e. p3-s1 and p4-s4, we obtained accuracy values statistically not different from chance level (p>0.05, binomial test, see Fig. 2c and Supplementary Fig. 1c for confusion matrices), i.e. 45.7% for p3-s1 ([36.8%, 54.7%], Confidence Interval p=0.05) and 52.3% for p4-s4 ([43.3%, 61.2%], Confidence Interval p=0.05). In the same animals, but with different electrode positioning, we achieved a high level of accuracy of 95.5% for p3-s2 ([90.5%, 98.31%], Confidence Interval p=0.05) and 75% for p4-s3 ([66.6%, 82.2%], Confidence Interval p=0.05, see Fig. 2c and Supplementary Fig. 1c for confusion matrices). Interestingly, in this scenario TIMEs and SELINEs performed similarly. Indeed, if we consider only the maximum accuracies achieved in each animal, thus neglecting possible confounding effects of electrode positioning, the TIME t1 in p2 achieved the maximum accuracy across animals in the case of baseline vs. BPC (Fig 2a), while SELINE s2 implanted in p3 performed better in the baseline vs. TVC, achieving a maximum accuracy in the animal p3-s2 (Fig. 2c). No significant differences between maximum accuracies were found for baseline vs. RRC (see Fig. 2b, p2-t2 vs. p4-s3 and p2-t2 vs p3-s2, p>0.05 Chi square test, Bonferroni correction).

The differences between two levels of RRC and TVC in p2 (from 10 to 15 respiratory cycles/min and from 400 to 500 ml, respectively) compared to p3 and p4 (from 10 to 20 respiratory cycles/min and from 400 to 800 ml, respectively) did not indicate strong differences in the decoding performances. In fact, if we compared the maximum accuracies obtained in a single animal (i.e., to neglect the effect of a possible dependency of electrode position) only in the case of baseline vs. TVC, the accuracy achieved in p2-t2 was lower than the maximum one observed in p3-s2 (Fig. 2c, p<0.05 Chi square test, Bonferroni correction), while no statistical differences were found (p>0.05 Chi square test, Bonferroni correction) in the other cases (Fig. 2b p2-t2 vs p3-s4, p2-t2 vs p4-s3, and Fig. 2c p2-t2 vs p4-s3).

Finally, a dependence on electrode position was also observed in pigs p4 and p3 for the comparisons of baseline vs. BPC (Fig. 2a) and baseline vs. RRC (Fig. 2c), respectively. In fact, the accuracy value for p4-s3 (96%, [92.5%, 98.1%] Confidence Interval p=0.05, Fig. 2a) was greater than p4-s4 (68.5%, [62.2%, 74.3%] Confidence Interval p=0.05, Fig. 2a) (p<0.001 Chi square test, Bonferroni correction). Moreover, in pig p3 implanted with SELINE s1 and s2, i.e. p3-s1 and p3-s2, we achieved a lower level of accuracy for p3-s1 (74%, [66.1%, 80.9%] Confidence Interval p=0.05, Fig. 2b) with respect to p3-s2 (93.2%, [87.5%, 96.8%] Confidence Interval p=0.05, Fig. 2b) (p<0.001 Chi square test, Bonferroni correction).

### Decoding the response to multiple functional challenges

We increased the complexity of the decoding task by producing two simultaneous functional challenges in different physiological systems, e.g. BPC plus RRC or TVC, or in the same system, namely RRC and TVC. As expected, the highest accuracy levels were achieved when alterations were simultaneously induced in different systems (Fig. 3a,b and Supplementary Fig. 2a,b).

**Fig. 3.**
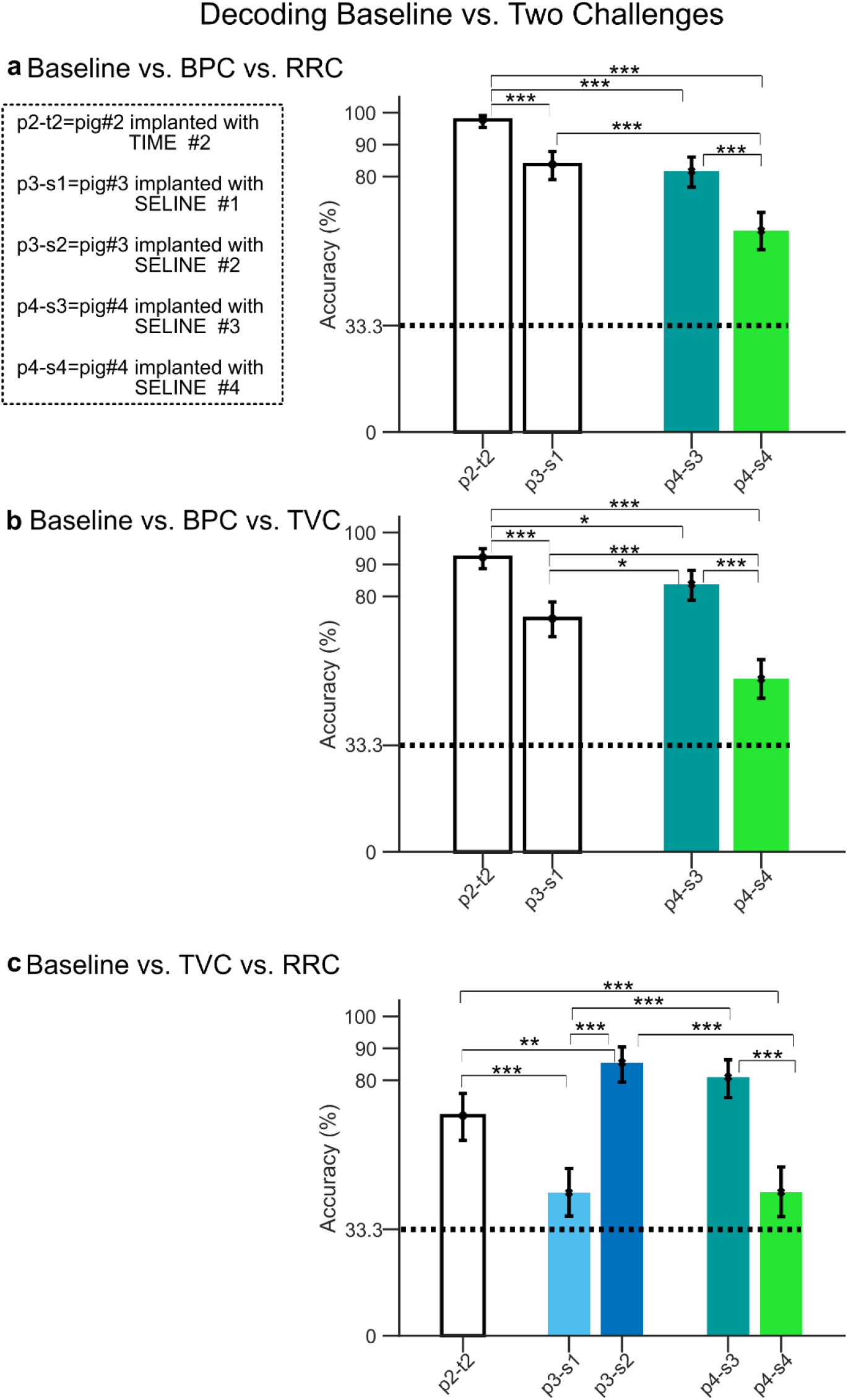
Decoding accuracy for baseline condition vs. two functional challenges. Accuracy level together with confidence intervals (error bars, Clopper-Pearson method) for the comparison between baseline and BPC vs RRC (panel a), BPC vs TVC (panel b), RRC vs TVC (panel c). For all panels, colored bars indicate the same animal implanted with two different electrodes. Dashed lines represent chance level. Statistical comparisons between animals were assessed by using a Chi square test Bonferroni corrected for multiple comparisons (*p<0.05,**p<0.01,***p<0.001).

The accuracy values for the BPC vs. RRC case were 81.6±7.1 % (n=4), as shown in Fig. 3a and Supplementary Fig. 2a for confusion matrices. Statistical differences between accuracy values of p4-s3 (81.7%, [76.7%, 86.1%] Confidence Interval p=0.05) and p4-s4 (81.7%, [57.1%, 68.8%] Confidence Interval p=0.05) indicate dependency on electrode position (p<0.001 Chi square test, Bonferroni corrected).

Dependency on electrode position was observed also for BPC vs TVC: in the animal implanted with s4, i.e. p4-s4, we found a statistically significant different accuracy level (54.2%, [48.1%, 60.2%] Confidence Interval p=0.05), than for p4-s3 (83.9%, [78.8%, 88.1%] Confidence Interval p=0.05) (p<0.001 Chi square test, Bonferroni corrected, see Fig. 3b). Overall, we obtained a mean over recordings 75.8±8.2% accuracy (n=4), as shown in Fig. 3b and Supplementary Fig. 2b for confusion matrices.

We were also able to reliably decode the two respiratory challenges RRC and TVC when occurring simultaneously. We found that in animals p3 and p4 the electrodes s2 and s3 achieved a high level of accuracy, 85.6% ([79.4%, 90.4%] Confidence Interval p=0.05) and 81% ([74.6%,86.4%] Confidence Interval p=0.05), respectively, as shown in Fig. 3c and Supplementary Fig. 2c. In contrast, in the same animals, the electrodes s1 and s4 implanted in different positions were only able to reach accuracy values equal to 44.8%([37.5%, 52.3%] Confidence Interval p=0.05) and 45%([37.2%, 52.8%] Confidence Interval p=0.05) for p3-s1 and p4-s4, respectively, as shown in Fig. 3c and Supplementary Fig. 2c. Moreover, in p1 implanted with TIME t2, we achieved accuracy values equal to 68.9% ([61.2%, 75.9%] Confidence Interval p=0.05). In this configuration, TIMEs seemed to perform better than SELINE in two cases, as the maximum accuracy for the cases BPC vs. RRC and BPC vs.

TVC were obtained by p2-t2 (Fig. 3a,b, p2-t2 vs p3-s1and p2-t2 vs. p4-s3, p<0.001 Chi square test, Bonferroni correction, respectively). However, the maximum accuracy for TVC vs. RRC was obtained by SELINE s2 implanted in p3, i.e. p3-s2 (Fig. 3c, p3-s2 vs p2-t2, p<0.01 Chi square test, Bonferroni correction).

Finally, we studied the most complex scenario of our experimental dataset in which all types of functional challenges (one cardiovascular and two respiratory) are decoded simultaneously. As discussed above, also in this case we found that decoding performances depended on electrode positioning within the same animal p4. The accuracy levels were 73.7% for p4-s3 ([68.5%, 78.5%] Confidence Interval p=0.05) and 45.3% for p4-s4 ([39.7%, 51%] Confidence Interval p=0.05), as shown in Fig. 4a and Fig 4c for confusion matrices. The highest accuracy among animals was obtained by TIME t2 implanted in p2, i.e. p2-t2, with a value equal to 87.3% ([83.3%, 90.6%] Confidence Interval p=0.05) (Fig 4a).

**Fig. 4.**
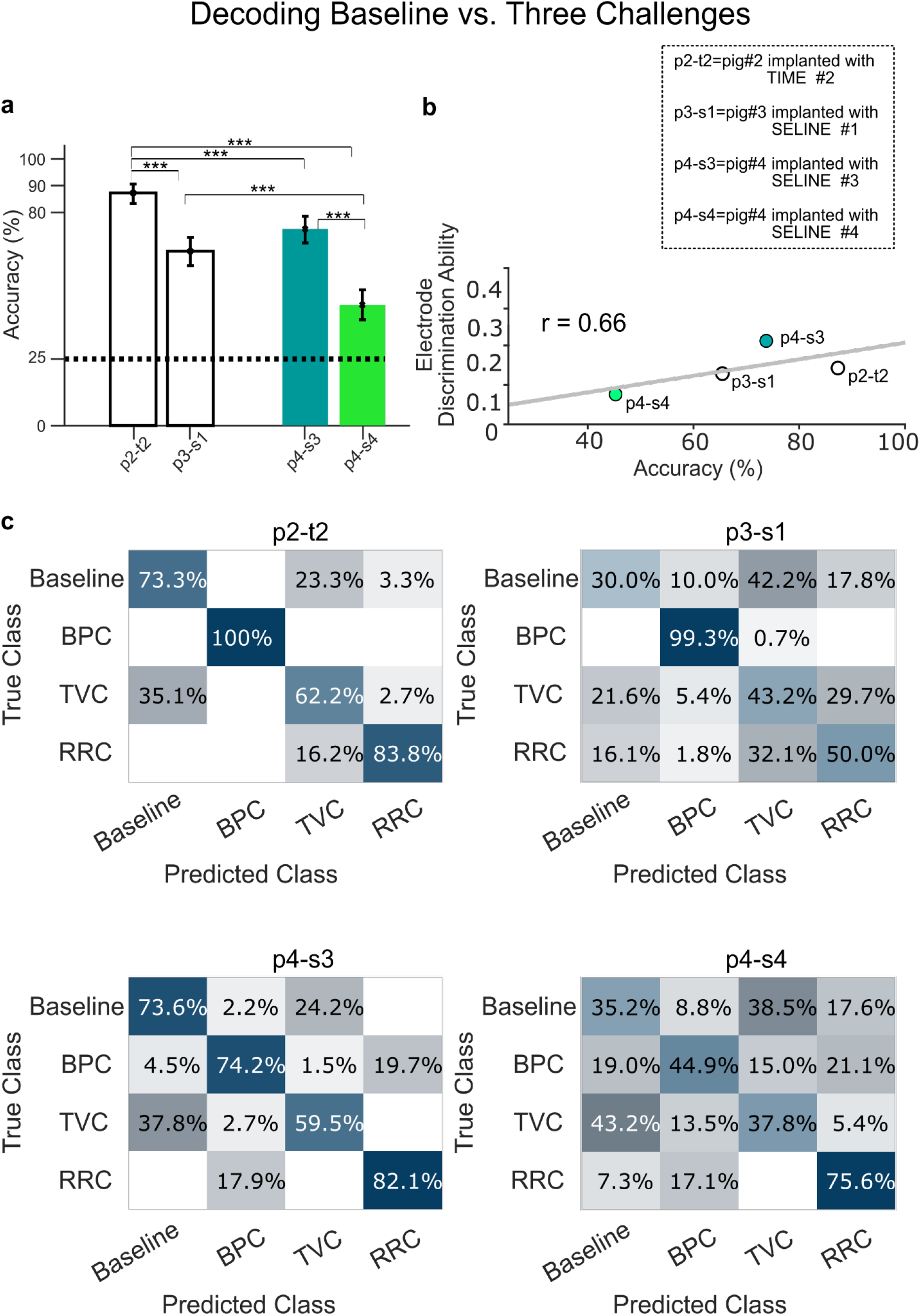
Decoding accuracy for baseline condition vs. 3 functional challenges and relation between electrode discrimination ability and decoding performances. **a** Accuracy level together with confidence intervals (error bars, Clopper-Pearson method) for the comparison between baseline and BPC, TVC and RRC. Colored bars indicate the same animal implanted with two different electrodes. Dashed lines represent chance levels. Statistical comparisons between animals were assessed by using a Chi square test Bonferroni corrected for multiple comparisons (*p<0.05, **p<0.01, ***p<0.001). **b** Linear relationship between electrode discrimination ability and accuracy level, linear correlation coefficient is reported. **c** Confusion matrices for all the recordings.

Furthermore, to better understand why different accuracy values were obtained in our datasets, we searched for a possible correlation between the electrode discrimination ability, i.e. the sensitivity of the recording sites to the specific challenges, and the decoding performance. Electrode discrimination ability (EDA) was assessed by calculating the percent activation of each recording site of the electrode with respect to the different functional challenges (see Methods for details and Supplementary Figures 3 and 4 for a graphical example). In this way, the ensemble of channels activations forms a discrimination vector for each functional challenge. Intuitively, the greater the difference between those discrimination vectors representing different functional challenges, the more discriminative is the electrode (Supplementary Fig.4b,c for distance matrices). We then calculated the linear correlation coefficient between EDA and accuracy (Fig. 4b) and found a correlation value, towards statistical significance, between electrode discrimination properties and decoding performance (Fig. 4b Pearson correlation coefficients for accuracy vs. EDA *r* = 0.66 and *n* = 4, *p* = 0.3).

### A hybrid model framework to map the functional spatial organization of VN fascicles

The site-related sensitivity to specific functional challenges indicated a possible functional spatial organization of VN fascicles, which we explored by combining the information obtained from histological analysis with simulations of electric potential field and discrimination properties of the electrodes.

The spatial relationship between VN fascicles and electrode active sites was determined by histological examination in one animal, p4 (see Methods for details). We reasonably assume that the overall nerve morphology in terms of fascicular structure and organization is constant along the implant site, so we morphologically modelled the SELINE active sites on the same 2D transverse section representation of the VN (Fig. 5 panels a, b, left and middle figures, for SELINEs s3 and s4 implanted in p4, respectively).

**Fig. 5.**
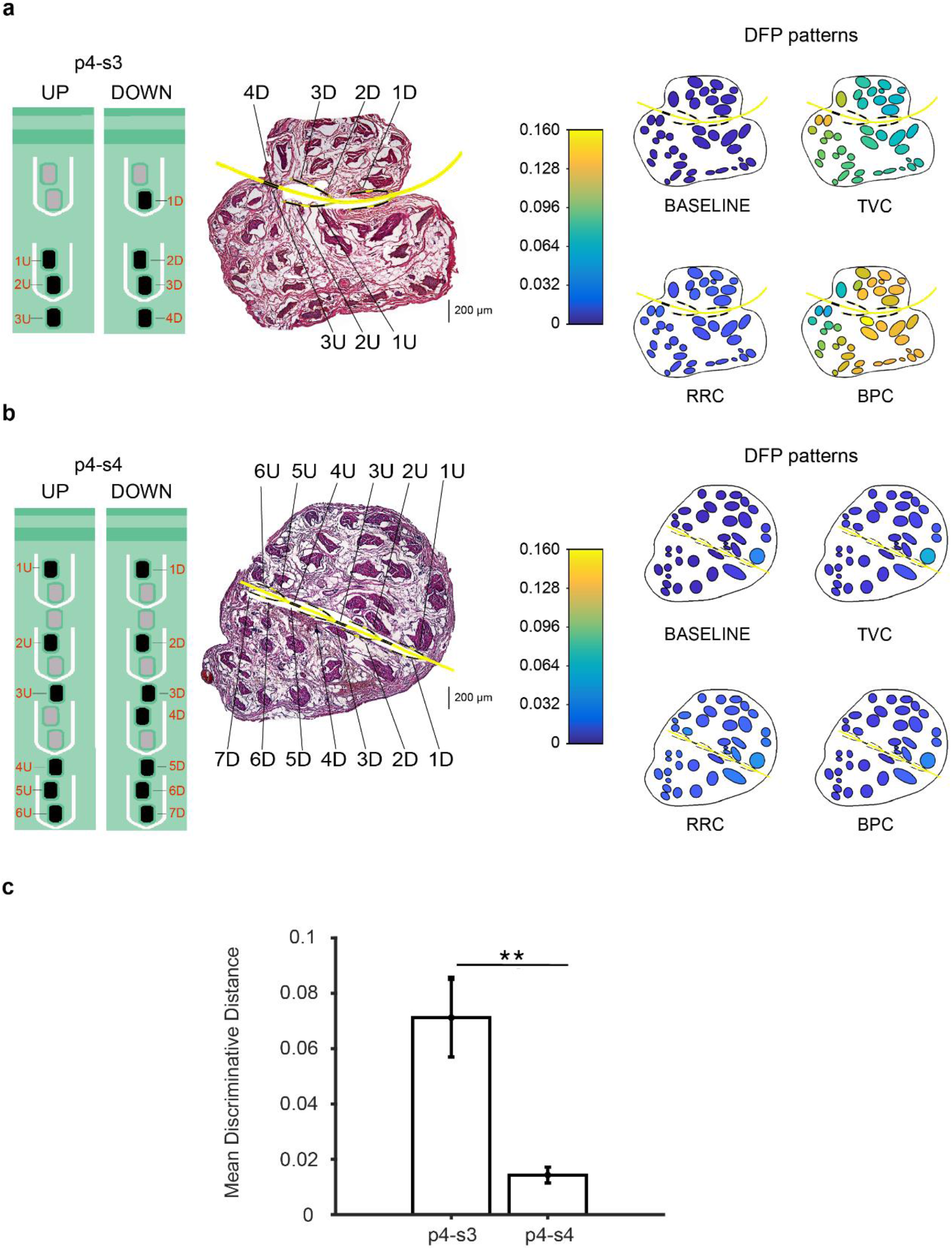
Schematic description of the two SELINE (s3 and s4) implantation in the VN of p4 and hybrid model simulations of fascicles activation at baseline and in response to functional challenges. **a,b** SELINE s3 and s4 (left panels a,b, respectively) are represented on a VN histological section (central panels a,b respectively). DFP patterns of activity during baseline and each functional challenge (left panels). **c** Mean Discriminative Distance between DFP patterns (p=0.003 unpaired t-test n=6, p>0.05 Kolmogorov-Smirnov).

Based on the histological analysis, we constructed a hybrid modeling framework^38,39,41^ to understand the spatial distribution of electrical potential fields in the nerve sections of p4-s3 and p4-s4 (see Methods for details). In Supplementary Figures 5 and 6 (panel b), isopotential electric field lines corresponding to each nerve section and 3 different recording sites are shown (p4-s3 and p4-s4, respectively). Using the Helmholtz reciprocity theorem^41^, these potential values quantify the recording capability of a given active site favored by the specific position of a fascicle located inside the nerve^42^. Since we were interested in the global electrical activity of fascicles, we employed a mean field approach by averaging the potential values within a given fascicle (Supplementary Figures 5 and 6 panel c, left) for nerve section p4-s3 and p4-s4, respectively.

To quantitatively characterize the association of a given fascicle with a specific functional challenge, we weighted the discrimination index of each active site with the mean field generated by that fascicle recorded from that active site. We then averaged data obtained from all recording sites and normalized the average to the total mean field, thus obtaining what we called Discriminative Field Potential (DFP, see Methods for details). In this way, we developed for the first time a quantitative measure characterizing how a fascicle is related to a given functional challenge. This measure could be considered the equivalent, for recording, of the selectivity index^20,38^ quantifying the stimulation ability on an electrode to elicit the activation of targeted fibres.

DFP patterns related to baseline condition and functional challenges were obtained for each nerve section p4-s3 and p4-s4, as shown in Figure 5 a,b, right panels, respectively, and Supplementary Figures 5 and 6 panel c bottom, respectively. Interestingly, for both nerve sections p4-s3 and p4-s4, the baseline condition exhibited a balanced DFP pattern where each fascicle is activated in the same manner. On the contrary, DFP magnitude was higher and differences between DFP patterns within each nerve were more pronounced in p4-s3 than in p4-s4 as shown in Figure 5 a,b, right panels, respectively, and Supplementary Figures 5 and 6 panel c bottom, respectively. We quantified them by calculating the root mean square level of the Euclidean distance among the DFP values of each fascicle for each possible comparison between functional challenges (mean discriminative distance). We found a statistically significant higher mean discriminative distance of the nerve in p4-s3 with respect to p4-s4 (p=0.003 unpaired t-test, n=6 all possible functional challenges combination). This was consistent with the higher effectiveness of decoding based on p4-s3 recordings.

## Discussion

Bioelectronic medicine may lead to revolutionary treatments for an ample variety of diseases. However, therapeutic neuromodulatory interventions must be very precise, both spatially, activating specific nerve fibres to avoid side effects, and temporally, i.e., operating in closed-loop to mimic the natural conditions. Therefore, it is necessary to utilize neural interfaces to selectively stimulate ANS nerves (spatial precision) and to decode signals triggered by specific functional changes, so that the stimulation is activated only when necessary (temporal precision) and subjected to a feedback control.

The present study provides the first evidence that signals recorded with intraneural electrodes in pig VN can be reliably used to detect functional changes in the cardiovascular and/or respiratory system.

We perturbed the homeostasis of one of both systems with functional challenges that were expected to enhance the activity of aortic baroreceptors and/or lung stretch receptors. Electrical signals departing from those receptors run along afferent fibres of the VN, carrying to the CNS information on blood pressure changes and lung inflation state^43^. Although induced with pharmacological and mechanical stimuli, these hemodynamic and respiratory alterations mimic very common pathophysiological conditions such as hypertension, tachypnea, and polypnea. Exploiting our previous experience with somatic nerves^31^ and differently from previous studies^3,21,23,24,32^ that employed spike sorting methods^23,24^ or extracted neural profiles correlated with specific physiological variables^3,21,32^, we considered the whole high frequency components of our signals processed with a novel advanced machine learning approach. This method proved successful in achieving high-level accuracy for the decoding of the functional challenges. In such regard, the pig VN is ideal for the development of clinically relevant technologies. The fascicular organization of pig VN is the closest to the complexity of human VN intraneural morphology compared to other species commonly utilized in experimental research^44^. For instance, other studies have shown the possibility to decode different functional challenges in the murine VN^23,24^, which, unfortunately, contains less fibres which are not subdivided in fascicles.

To the best of our knowledge, prior studies in pig VN^3,21,32^ did not demonstrate the possibility to decode multiple functional challenges. While cuff^3,21^ and intraneural^32^ electrodes in pigs proved effective in extrapolating neural markers of blood pressure and respiratory activity, no decoding analysis for the identification of mixed functional challenges was attempted. Moreover, the recordings with cuff electrodes^3,21^ were not obtained during the same experimental session.

In this study we exploited intraneural (TIMEs and SELINEs) electrodes, which are conceived to be transversally or obliquely inserted into the nerve, allowing spatially selective stimulation and recording from different fascicles innervating distinct peripheral targets^45,46^.Our results show that the quality of decoding performances depends on the electrode position. This could be due to the different functional role of fascicles adjacent to the recording sites and prompts the hypothesis of a specific spatial segregation of vagal fibres traversed by specific signals. Therefore, to map a possible spatial functional organization of VN fascicles, we employed a hybrid modeling framework based on histological analysis combined with electrodes discrimination ability properties. We assumed that the discrimination of a given functional challenge in a specific recording site is higher when the local fibres are activated by that specific stimulus and the local recording capability is high. Based on this assumption, we developed a novel quantitative measure called Discriminative Field Potential (DFP), obtaining distinct spatial configurations of discriminative patterns generated by fascicles during the various functional challenges. It is important to note that the good reliability of DFP patterns resulted from the use of intraneural electrodes, which potentially provide a more accurate recording of the fascicles neural activity compared to epineural electrodes.

Moreover, placing a large number of active sites along the intraneural implant would augment the interfaces with fascicles, thus reducing the number of implanted electrodes^29^. The present findings strongly suggest that multi-contact electrodes positioning is of crucial importance for bioelectronic applications in a complex nerve such as the VN. In our opinion, the development of an anatomical and electrophysiological in silico model of the VN would be extremely useful to guide electrodes implantation and positioning and to overcome the present limitations. In this regard, the pig VN represents an excellent model, given its marked fasciculation, which likely reflects an equivalent high spatial segregation of fibres innervating different peripheral sites.

### Limitations and future directions

The present study was performed in an acute experimental preparation and it is known that anesthesia exerts important effects on neural activity and other physiological systems. For this reason, our next step will consist of validating the implantation of intraneural electrodes for VN chronic recordings in non-anesthetized animals. Moreover, our results seem to indicate that TIMEs could perform better than SELINE, but more experiments are necessary to confirm these results. Finally, another important step will be its implementation and validation of online signal analysis, necessary for closed-loop VNS protocols.

### Conclusions

The present results are an important step towards more precise neuromodulation protocols based on the knowledge of specific patterns of neural activations recordable in VN under physiological and pathological conditions. Advanced closed-loop and spatially selective VNS could improve the treatment of pathological conditions by selectively activating functionally specialized fibre fascicles, thus preventing the numerous side-effects caused by the current technology.

The importance of understanding VN signals extends beyond VNS and closed-loop applications. Being the biggest peripheral crossroad of signals between the CNS and visceral organs, a sensitive and reliable technology for detection and decoding of VN activity could serve for the diagnosis of pathophysiological conditions otherwise difficult to detect. Deciphering the “vagal language” will also help clarifying the mechanisms by which VNS exerts its curative effects and gaining deeper insights in the VN regulation of specific physiological processes.

## Materials and methods

### Animals

This study was conducted in four castrated male farm pigs (*Sus Scrofa Domesticus*) 3-4-month old and weighing 28-30 kg. The animals were housed in the vivarium at room temperature (24°C) and fasted overnight before anesthesia with free access to water. All animal handling and experimental procedures were performed according to European Community guidelines (EC Council Directive 2010/63) and the Italian legislation on animal experimentation (Decreto Legislativo D.Lgs 26/2014).

### Anesthesia and monitoring of physiological parameters

On the day of the experiment, pigs were anesthetized and prepared for surgery as previously described by us^47^. They were sedated with a cocktail of 4 mg/kg tiletamine hydrochloride and 4 mg/kg zolazepam hydrochloride injected intramuscularly, intubated and mechanically ventilated. The respiratory frequency was fixed to 10 respiratory cycles per minute and the tidal volume (TV) to 400 ml. The combination of these respiratory parameters resulted in PaO_2_ > 100 mmHg, PaCO_2_ < 40 mmHg and arterial blood pH in the range of 7.4-7.45. Arterial blood gases analysis was repeated at least every 30 min. A pulse oximeter was applied on the tongue to continuously monitor arterial oxygen saturation, which was stably above 96% for the duration of the experiment. Inhalatory anesthesia was maintained with a mixture of 1-1,5% isoflurane dissolved in 79% air and 21% oxygen. The respiratory pressure was recorded by connecting the airflow of the ventilator to a pressure transducer. Electrocardiogram (ECG), heart rate, and arterial blood pressure were constantly monitored. Aortic blood pressure was recorded using a solid-state pressure transducer catheter (Millar, SPR-100) inserted in the left femoral artery with the tip positioned in the abdominal aorta. This catheter was also used to withdraw arterial blood for blood gas analysis. Glycemia was checked every 30 min and before each recording session to assess the stability of plasma glucose levels above 100 mg/100 ml. The main ear vein was cannulated for the intravenous administration of all the drugs.

### Surgical preparation and electrode implantations

The 4 pigs were named p1, p2, p3 and p4. To isolate the cervical VN, a midline cervical incision was made from the level of the larynx to the sternum, as previously described by us^48^. After identifying the neurovascular bundle of the neck (common carotid artery, internal jugular vein and VN), the left (n = 3) and right (n = 1) cervical VNs were delicately separated from the common carotid artery by blunt dissection, 3-4 cm above and below the cricoid cartilage. The sympathetic trunk, attached dorsally to the VN, was gently detached and pulled apart. In animals p1 and p2, the left VN was implanted in correspondence of the cricoid cartilage with a transverse intrafascicular multichannel electrode (TIME) endowed with 10 active sites, following the same procedures described by Badia et al^49^. In p3 and p4, the left and right VNs, respectively, were implanted with 2 self-opening intraneural peripheral interfaces (SELINEs) endowed with 7-16 active sites. SELINEs implantation followed the same procedures described by Cutrone et al^46^. All the intraneural electrodes were inserted obliquely, forming a 45° angle with the longitudinal axis of the nerve, except for 2 SELINEs, one in p3 and one in p4, that were inserted along the transverse axis. In all the experiments, the ground electrode was inserted under the skin of the left elbow of the animal.

### Tissue isolation and histology

At the end of the experiment, the anesthetized pigs were euthanized by an intravenous injection of saturated KCl solution. Intraneural electrodes were left in place and the VN was cut 3 cm above and below the insertion site and gently placed on a dedicated support to avoid twisting. The VN samples were then fixed in 4% paraformaldehyde for 18-20 hours, rinsed in PBS, dehydrated in a series of progressively more concentrated solutions of ethanol/xylol and finally embedded in paraffin wax. Transverse sections (10 μm thick) were cut, deparaffinized, rehydrated, processed for routine hematoxylin and eosin and mounted in silane-coated slides. Images were captured with 5x magnification by a Leica DMRB microscope equipped with the DFC480 digital camera (Leica Microsystems, Cambridge, UK). Sections at the level of the implanted electrodes were aligned and manually segmented to study the electrode–nerve interaction. The nerve was cross-sectioned for the entire length of the implants, with the two SELINEs fixed in site, and stained with hematoxylin and eosin. Light microscopy observations of VN sections showed the fascicular structure of the nerve and holes caused by the electrode insertion, with SELINE polyimide strips preserved in a few slices. No macroscopic and microscopic signs of hemorrhages were found.

### In vivo recording of VN activity

Neural recordings started at least 30 min after electrode implantation to allow for stabilization of the nerve and of physiological parameters. The neural signals were first acquired over 5 minutes at baseline, followed by recording sessions in which physiological parameters were altered (functional challenges). The baseline condition was defined as the time BP and heart rate were found stable for at least 30 min after completion of electrodes implantation. At baseline, RR and TV were fixed at 10 respiratory cycles per minute and 400 ml, respectively. The functional challenges consisted of 1) increase in mean arterial BP up to 150% of the baseline value obtained with an intravenous infusion of Ang II (Sigma Aldrich), a vasopressor with rapidly reversible effects^50^, at 80 ng/kg/min 2) increase in TV from 400 to 500 in p2 and 800 ml in p3 and p4 for 2 minutes and 3) increase in RR from 10 to 15 in p2 and 20 in p3 and p4 respiratory cycles/min. These challenges were produced in random order and each one was followed by a recovery period of at least five minutes to let the physiological parameters return to baseline values. These functional challenges and the related physiological parameters affected were in the same range of the ones used by Sevcencu et al.^21^ and proved suitable for neural VN recordings in anesthetized pigs. In addition, they did not interfere significantly with the stability of other variables like BP, heart rate or blood gases partial pressures. Glycemia and blood gases were checked before and after the completion of each functional challenge.

### Data acquisition and decoding algorithm

VN raw multichannel signals were acquired at a frequency sampling of 24.4 kHz, high pass filtered at 5 Hz and digitally amplified (RZ5D BioAmp Processor, Pz5, Tucker-Davis Technologies Inc., TDT, USA). Each channel of the raw recordings was segmented by using a 1 sec-temporal window and rescaled to zero mean and unitary standard deviation. Discrete wavelet transform^33,34^, at a maximum level equal to 4, was applied on the segmented portions of each channel by using a symlet 7 wavelet function^13,33,34^. Approximation coefficients relative to frequencies ≲ 1500 Hz were discarded in the subsequent analysis. Principal component analysis on the multivariate wavelet details was applied independently on two different scales^35^ relative to frequency ranges of ≅1500-3000 Hz and 3000-6000 Hz, respectively. The first three principal components were retained to build a feature vector for the classification algorithm. The entire dataset was randomly split in 70 % for the train set and the remaining 30% was used as test set. Ensemble of classifiers were built using a classification tree^36,37^. To deal with class imbalance problems, random undersampling techniques combined with boosting algorithms were adopted (RUSBoost)^37,51^. To achieve higher ensemble accuracy, we set the maximal depth of the decision tree equal to the number of the element in the train set.

We set the learning rate and the number of cycles to 0.1 and 1000, respectively, in order to achieve higher accuracy as well.

Classification performance was assessed on the test set by calculating accuracy of the corresponding confusion matrices. The accuracy level was calculated by considering the number of correct predictions divided by the total number of elements in the test set. Confidence intervals for accuracy values were computed by using Clopper-Pearson method. Chance level was estimated as the proportion of the class with the majority of elements with respect to the total number of elements.

Data were analyzed off-line in MATLAB (The MathWorks, Inc.).

### Discrimination analysis and decoding-discrimination relationship

Similarly to previous studies^20^, we built a discrimination channel index to understand the discrimination ability of the electrodes and consequentially a possible relation with decoding performances. To this aim, we calculated the percentage of activation for each channel relative to the different functional challenges as described below.

For each challenge, each signal was reconstructed using the wavelet details at the same scale previously used for the decoding algorithm:

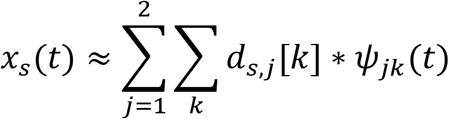

where *d_s,j_*[*k*] are the wavelet details at scale *j* of the recording site *s*, and *ψ_jk_*(*t*) is the wavelet function (see Supplementary Figure 3a,b for a graphical example). The reconstructed signals from each recording sites (*X* = {*x_s_*}, *i* = 1,…, *S*) were segmented using a temporal window of T=1 sec and rescaled to zero mean and unitary standard deviation obtaining a number of signal’s blocks *N_block_*, i.e. *X* = (*X*(1),…, *X*(*N_block_*)). For each block of the signals, we calculated the first three principal component coefficients (loadings), i.e. *P^i_pca_^*(*i_block_*) with *i_pca_* = 1,2,3 and *i_b1ock_* = 1,…, *N_block_*. We thus identified for each principal component the higher outliers relative to loadings’ absolute values. Outliers were identified using a threshold of three scaled mean absolute deviation from the median (see Supplementary Figure 3c, right panel for a graphical example). When an outlier was identified in a portion of the signal, a value equal to unity was assigned to the corresponding channel (‘channel activated’) and zero otherwise, i.e. we defined an indicator for each channel s and block *i_block_*:

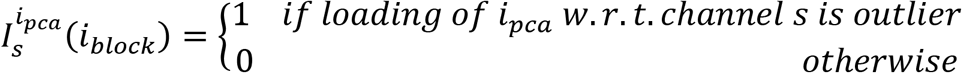

For each channel, the resulting percentage of activation relative to a principal component 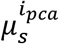, is equal to the number of times in which the channel was activated divided by the total number of portions of the signal, i.e.

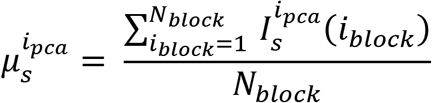

The percentage of activation of each channel was quantified as the mean over the percentage of activation of the channel in each principal component (see Supplementary Figure 3c left panel) 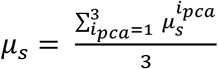.

Similarly to Raspopovic et. al^20^, given the percentage of activation for each channel, we defined a discrimination index for each channel as the difference between the percentage of activation of the considered channel and the mean percentage of activation over the remaining channels, i.e.

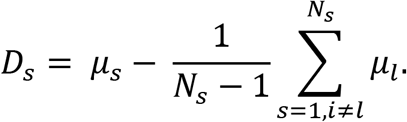

Thus, discrimination vectors with entries made of channels’ discrimination indices were obtained among the different functional challenges, i.e. *D_c_* = {*D_c,s_*} with *c* = {*Baseline, BPC, TVC, RRC*} and *s* = 1,…, *S*. The difference between two generic discrimination vectors was obtained by considering the root mean square level of the Euclidean distance (see Supplementary Figure 4a).

Electrode discrimination ability was computed by considering the mean over all the six possible combination distances between the distinct discrimination vectors (see Supplementary Figure 4a). Finally, a linear correlation coefficient was calculated between the electrode discrimination ability and the decoding performance (accuracy) and the least-squares line was plotted.

### Hybrid modelling of the recording process and Discriminative Field Potentials

To understand the distribution of the electrical field potential within a nerve, we employed the hybrid modeling framework as described in^38,39,41^. Histological sections were manually segmented by defining the contours of the epineural compartment (the whole nerve section contour) and of the fascicles. Such contours were imported in MATLAB as closed polylines. Fascicle polylines were substituted with ellipses having the same surface area as the fascicles, center of the fascicle centroid, axis length ratio and orientation were deduced from the minimum (area) bounding rectangle of each fascicle. The perineurial sheath for each fascicle was defined as a layer with width equal to 0.03 times the effective diameter (diameter of the equivalent circle) of the fascicle^52^. A 3D extrusion of the given nerve section was defined in COMSOL (through MATLAB LiveLink for COMSOL) and it was included in a saline bath whose external surface was grounded (simulation of zero potential at infinity^39^).

A point current source was defined corresponding to each electrode active site center. To obtain the potential field on the nerve section (up to a multiplicative scaling factor, i.e. linearity assumption), we simulated the stimulation from all given sites with an adimensional current (one at the time and unitary). Indeed, thanks to the Helmholtz reciprocity theorem, the value of the potential field due to a unitary current injected by an active site corresponds to the impedance that relates the intensity of a current source with the resulting potential field recorded at the site surface. This means that a higher electrical field in a point in space P leads to a higher amplitude of the recorded electrical activity of a fibre located in P^42^ Given the knowledge of the electric field potential generated from a fascicle and the corresponding recording capability of a given recording site, we wanted to link this information to the discrimination properties of the recording site. To this aim, we defined a measure, that we called Discriminative Field Potential (DFP), by weighting the recording capabilities of a recording site *s* from a given fascicles *F* together with the discrimination properties of the recording site *s* related to a functional challenge *c*, i.e.

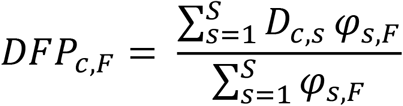

where *D_c,s_* is the discrimination ability to a functional challenge *c*and recording site s, and *φ_s,F_* is the mean field inside a fascicle *F* recorded from a site *s* (calculated by averaging isopotential values of the potential field inside the fascicle). Since the *φ_s,F_* measures how a site *s* is capable of recording from a given fascicle, and *D_c,s_* measures how site *s* records better the activity related to a functional challenge *c* with respect to the other sites, then the *DFP_c,F_* is a quantity measuring how a fascicle is related to a given functional challenge.

Similarly to the calculation of electrode discrimination ability, to quantify the discrimination ability of the whole nerve section, we computed the root mean square level of the Euclidean distance of the *DFP* values in the fascicle space. We thus calculated the mean discrimination distance by considering the mean over all the six possible combinations of functional challenges.

### Statistical analysis

Unless otherwise stated, data are expressed as mean ± s.e.m. Statistical significance between accuracies were assessed by means of Chi square test Bonferroni corrected for multiple comparisons. Confidence intervals for the accuracy values were calculated using the Clopper-Pearson method. Statistical significance of accuracy values with respect to chance level was assessed by using the binomial test. Statistical significance between the discrimination ability of nerve sections was assessed by using two-tailed unpaired t-test at a significance threshold equal to 0.05. To test for normality of data distribution a Kolmogorov-Smirnov test was used on each population of data. Statistical analysis was performed with MATLAB (The MathWorks, Inc.).

## Supplementary Information

**Supplementary Figure 1.**
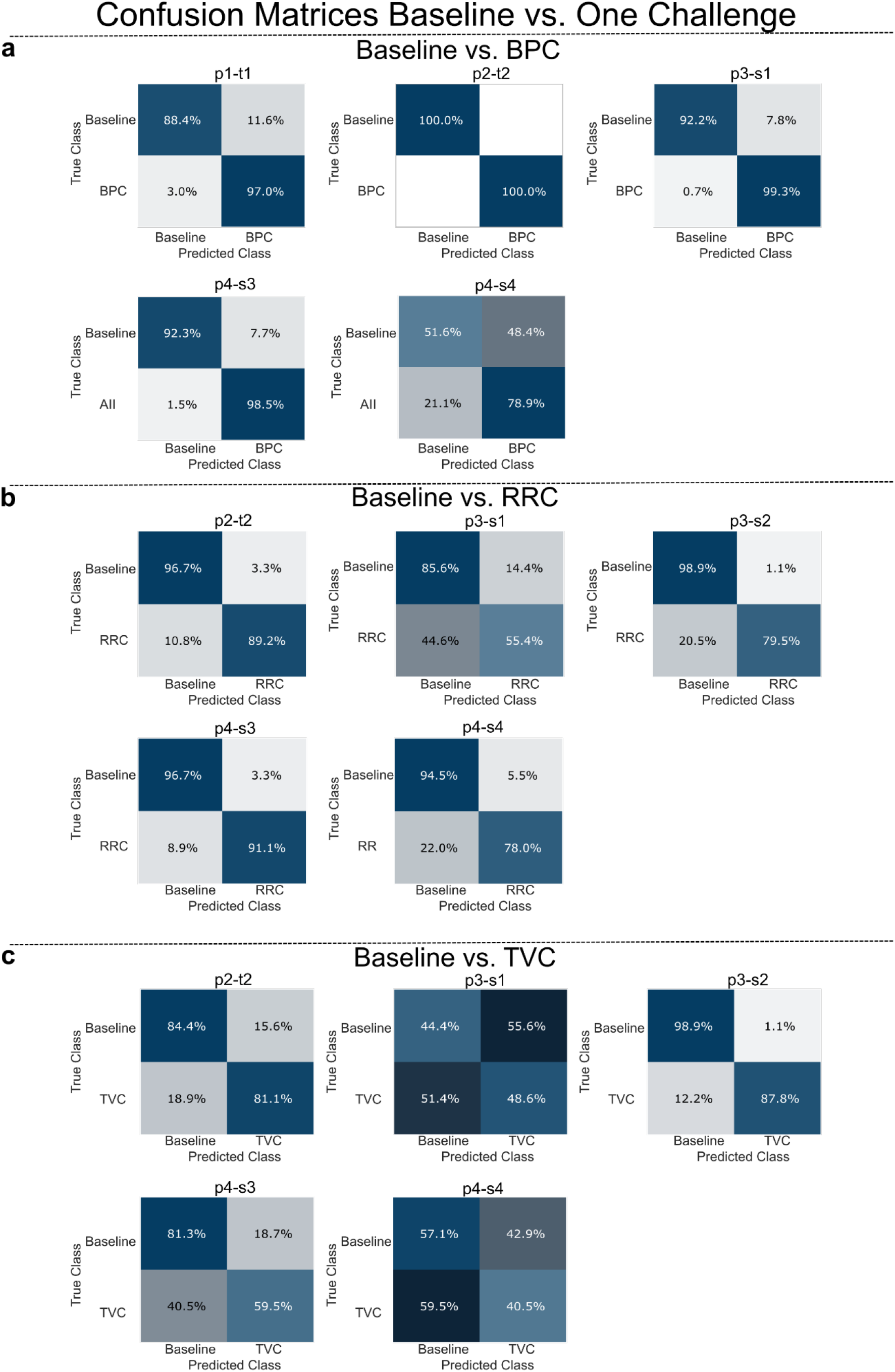
Confusion matrices between baseline condition against one functional challenge a,b,c. Comparison between baseline and BPC (panel a), RRC (panel b), TVC (panel c).

**Supplementary Figure 2.**
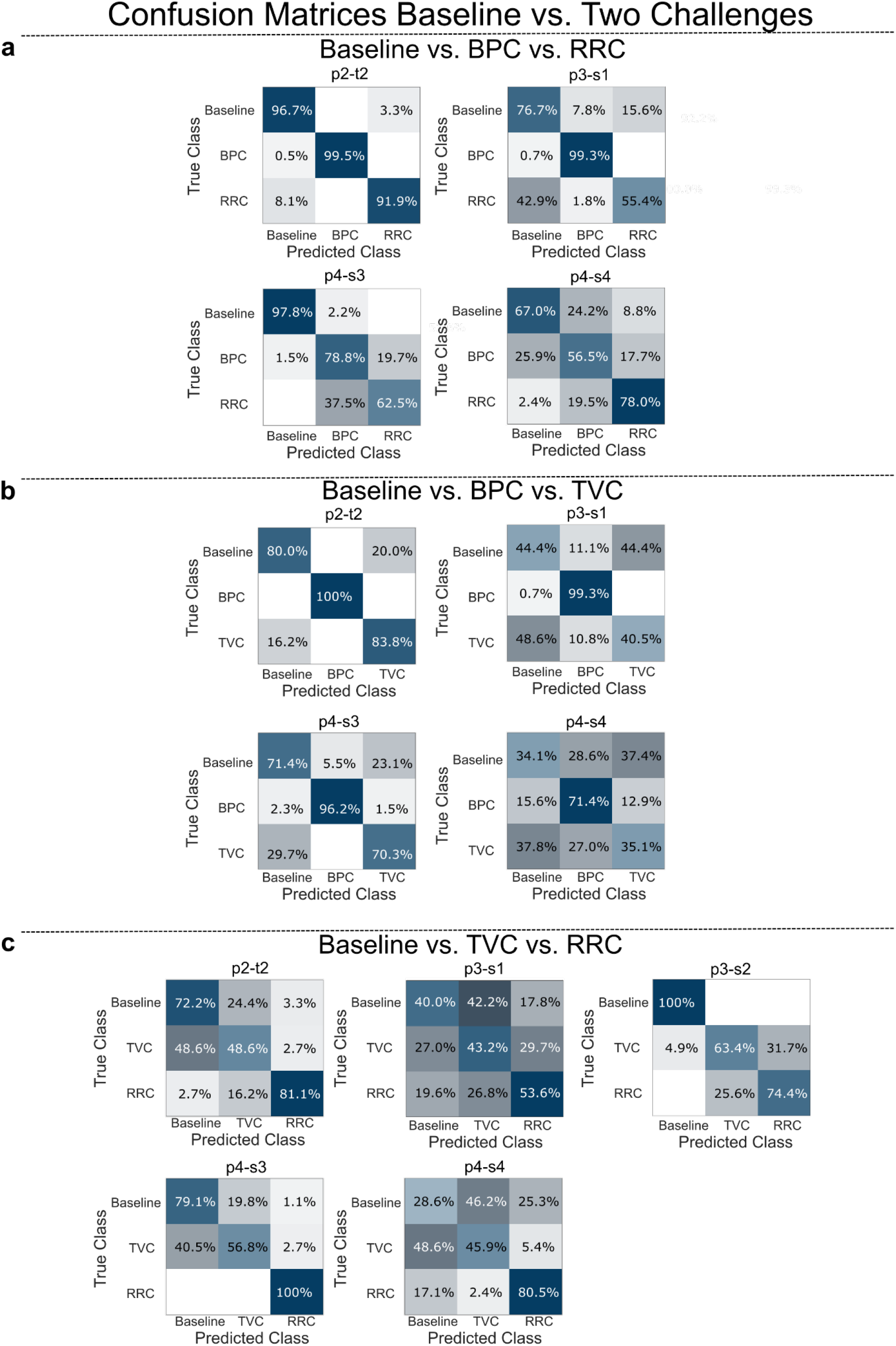
Confusion matrices between baseline condition against two functional challenges a,b,c. Comparison between baseline and BPC-RRC (panel a), BPC-TVC (panel b), RRC-TVC (panel c).

**Supplementary Figure 3.**
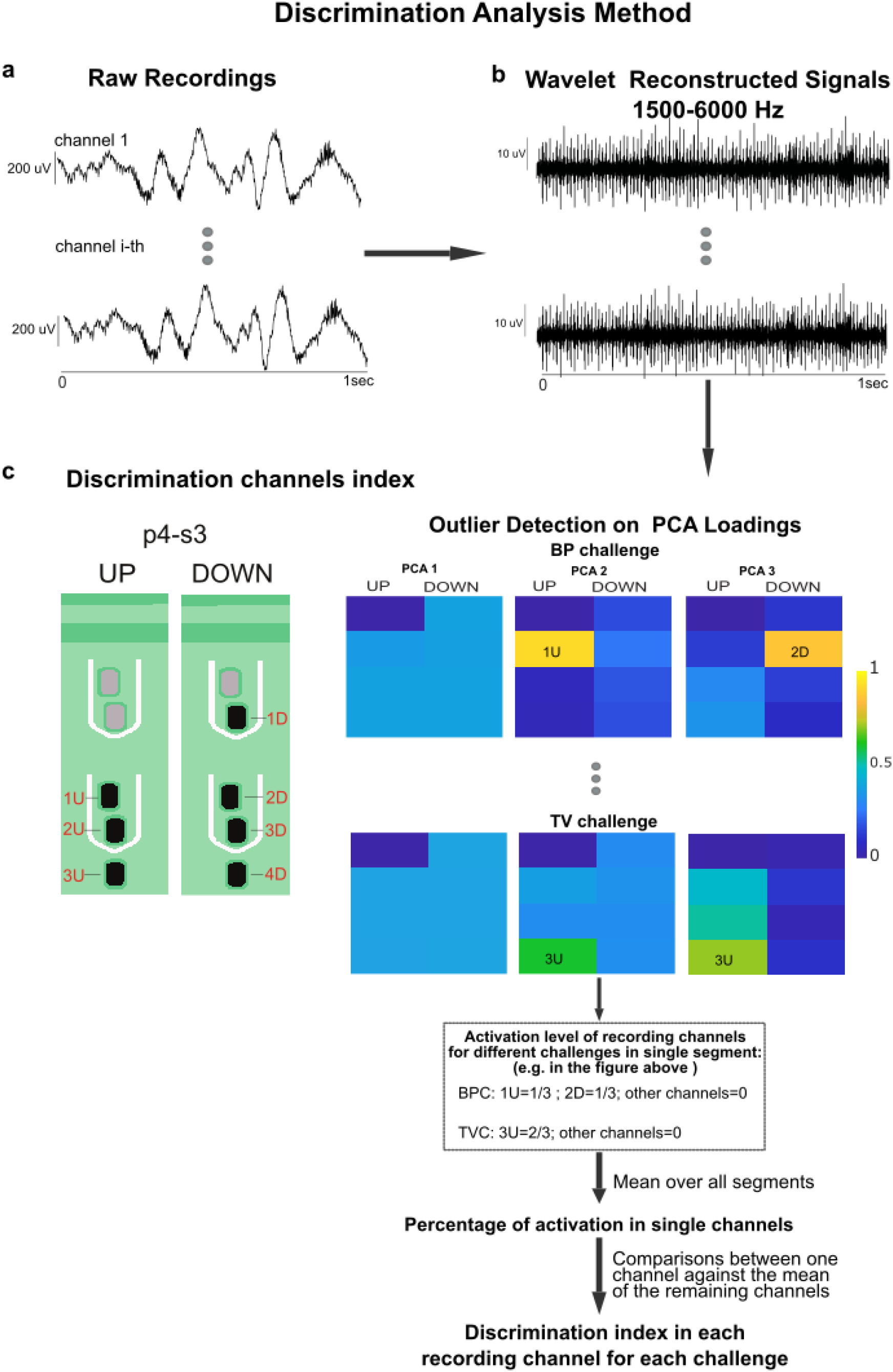
Discrimination analysis method. **a** Raw recording. **b** Signal in the frequency band 1500-6000 Hz reconstructed throughout wavelet details used in the decoding algorithm (segmented in a temporal window of 1 sec). **c** Estimation of channels discrimination index. Right panel, outlier detection on PCA loadings for the different functional challenges. Left panel, schematic representation of the active sites in the electrode p4-s3 and percentage of activation level estimation in single channels.

**Supplementary Figure 4.**
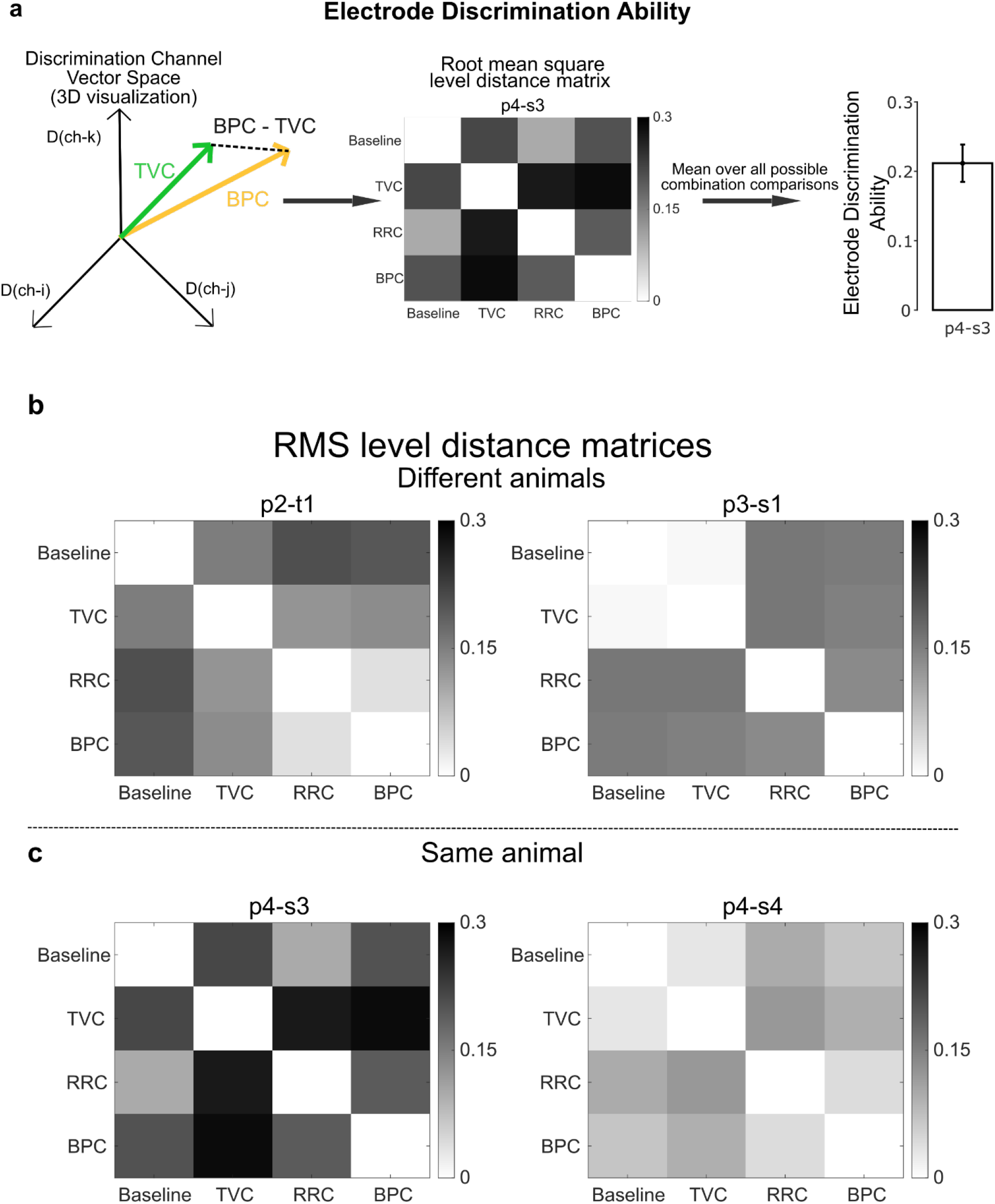
Estimation of electrode discrimination ability and RMS level distance matrices examples. **a** Estimation of electrode discrimination ability. Left panel, 3D graphical visualization example of discrimination index channel vectors for TVC and BPC challenges (green and yellow vectors, respectively). Middle panel, root mean square level distance matrix. Right panel, estimation of electrode discrimination ability by averaging over all possible (n=6) combination comparison of functional challenge. b,c RMS level distance matrices in the electrode discrimination space for the comparison among baseline condition and the functional challenges (TVC, RRC, BPC) in two electrodes placed in different animals (p2-t2 and p3-s1) and two electrodes in the same animal (p4-s3 and p4-s4), respectively.

**Supplementary Figure 5.**
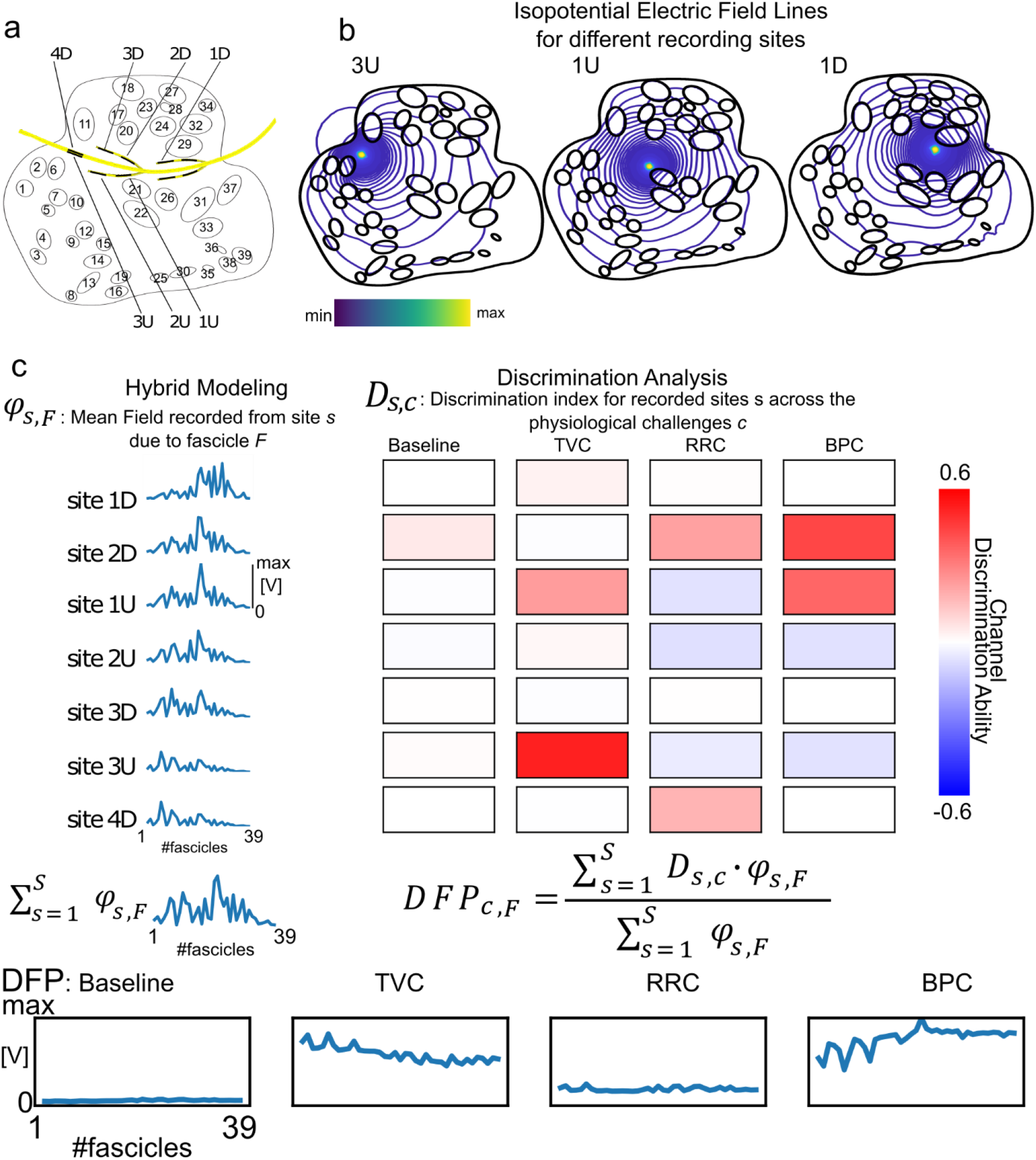
Discriminative Field Potential (DFP) estimation in pig p4e1 and p4e2, respectively. **a** Nerve cross-section and implanted electrode. The intraneural fascicles numerical are labeled with numbers i=1,…,N_fascicles_. **b** Isopotential Electric field lines for three different recording sites. **c** DFP estimation procedure. Left panel, fascicle mean field for each site. Right panel, selectivity index for each site across the different functional challenges. Bottom panel, DFP values in each fascicle across the different functional challenges.

**Supplementary Figure 6.**
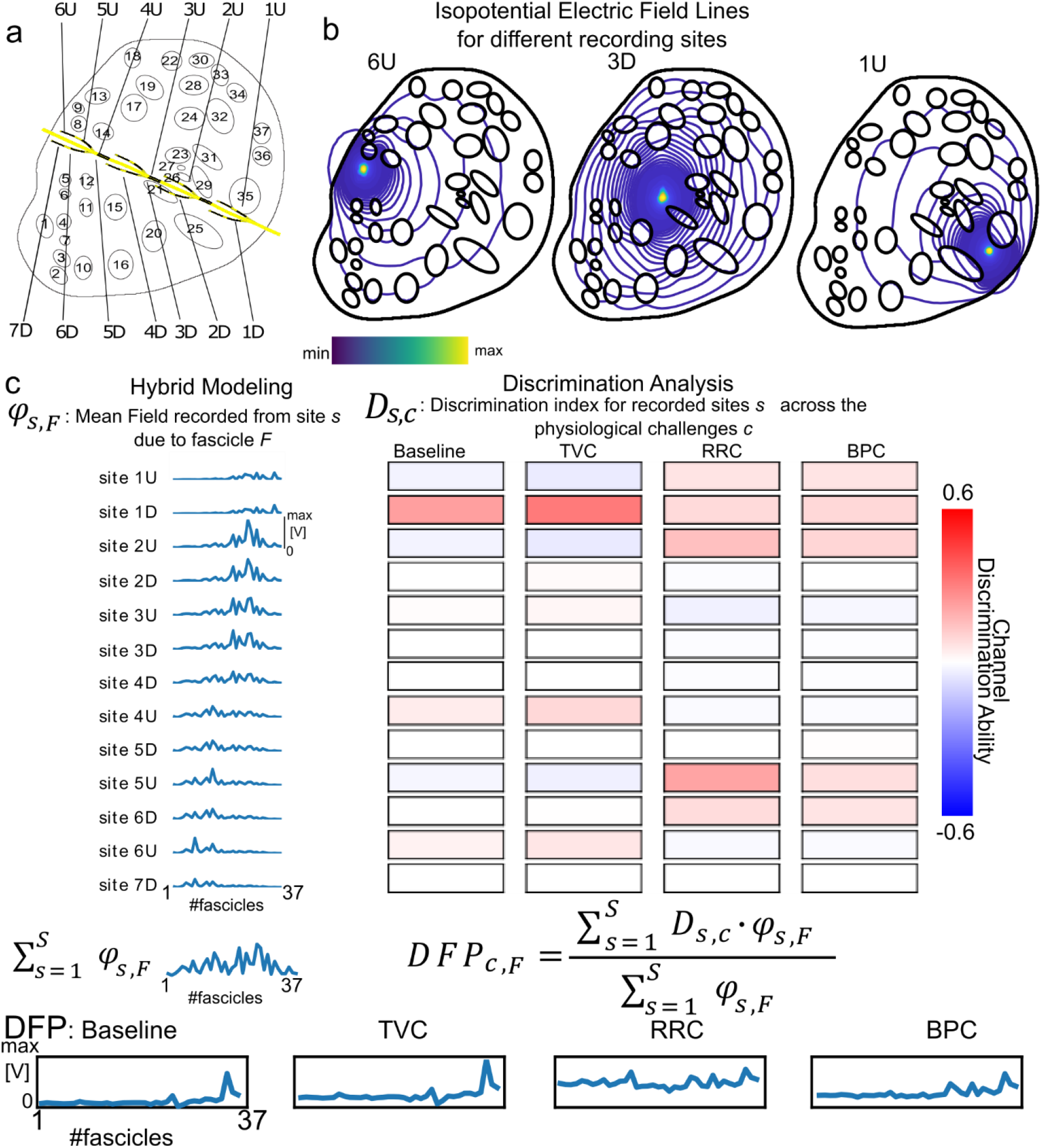
Discriminative Field Potential (DFP) estimation in pig p4e1 and p4e2, respectively. **a** Nerve cross-section and implanted electrode. The intraneural fascicles numerical are labeled with numbers i=1,…,N_fascicles_. **b** Isopotential Electric field lines for three different recording sites. **c** DFP estimation procedure. Left panel, fascicle mean field for each site. Right panel, selectivity index for each site across the different functional challenges. Bottom panel, DFP values in each fascicle across the different functional challenges.

